# Taxonomic Treatment of *Blighia* K.D.Koenig occurring in Ghana

**DOI:** 10.1101/2024.04.30.591818

**Authors:** Samuel Larbi, Hanan Abdul Fatah Lateef, Bismark Anum, Benjamin Darko Williams

## Abstract

*Blighia*, a genus consisting of trees in the Sapindaceae, is found in sub-Saharan Africa and other parts of the world. It is characterized by three species globally: *B. unijugata* Baker, *B. sapida* K.D. Koenig, and *B. welwitschii* (Hiern) Radlk but *B. unijugata* is absent in Ghana. Research on *Blighia* in West Africa has been primarily conducted in Nigeria, Togo, and Benin. However, the taxonomy of the genus is lacking in West Africa, with the Flora of West Tropical Africa being the only authoritative literature. A revision of this genus in Ghana is needed due to the monotonous concentration of research on *B. sapida* and the knowledge gap from Hutchinson & Dalziel. This paper seeks to update the taxonomy of members in *Blighia* occurring in Ghana.

The study collected specimens of *Blighia sapida* and *B. welwitschii* from 15 localities across Ghana. Fossil records, voucher specimens, and digital collections obtained from the Ghana Herbarium and GBIF respectively, were also examined. Standard georeferencing software was used to generate distribution maps.

*B. sapida* produces oblanceolate or obovate leaves with rounded or emarginate apices and acute bases. They are mostly distributed along the coastal belt of the country while *B. welwitschii* produces lanceolate leaves with bluntly attenuate or acute apices and cuneate bases. They are however distributed mainly in the middle belt of the country.

## INTRODUCTION

*Blighia* is a member of the Order Sapindales and the family Sapindaceae. The generic name *Blighia* is traced to Captain William Blighia; who moved a nameless tree from Jamaica to the Kew Gardens in England in 1793 (Rashford, 2001). It is widely spread across sub-Saharan Africa and in different parts of the world like the Caribbean, North and South America, and Europe. There are three accepted species in the genus: *Blighia unijugata* Baker, *Blighia sapida* K.D. Koenig, and *Blighia welwitschii* (Hiern) Radlk. (Sinmisola et al., 2019). However, in Ghana, there are two species: *B. sapida* and *B. welwitschii*.

*B. unijugata*, an indigenous tree in West Africa, is known for its unique single-leaflet leaves and is used as a shade tree. Its fruits are used as fish poison, and its bark can be used for building and treating illness (Sofidiya et al., 2012; Hyde et al., 2024). *B. sapida*, is a torrid perennial tree with wide, pinnate leaves and distinctive green foliage. It is distributed across West Africa, including Ghana, Liberia, Togo, Senegal, Benin, Cameroon, and Nigeria. Its red-yellowish fruits and oil are used for food and soap making (Omobuwajo et al., 2000; Aloko et al., 2017; Sinmisola et al., 2019). *B. welwitschii* is an evergreen tree with a dense crown. It is spread across Tropical Africa and used for food, medicine, and wood, with unique leaves composed of multiple leaflets. However, its bark, young leaves, fruits, and seeds can be toxic to fish in Sierra Leone (Burkill, 2000; Obeng, 2010; Ken Fern, 2024).

Greater portion of the research on *Blighia* in West Africa has been spearheaded by Nigerians (Omobuwajo et al., 2000; Abolaji et al., 2007; Atolani et al., 2009; Sinmisola et al., 2019) and other neighboring countries like Togo (Alfa et al., 2016; Kola et al., 2020; Nabede et al., 2022), Benin (Ekué et al., 2008; Ekué et al., 2011; Kakpo et al., 2020; Ndiaye et al., 2022). The only authoritative research on the genus in Ghana was conducted by Quartey et al. (2022). The contributions of the other West African countries have helped advance the knowledge of *Blighia* in the fields of medicine, agriculture, nutrition, taxonomy, and agroforestry. However, the taxonomy of the genus is lacking in West Africa. The Flora of West Tropical Africa is still the only authoritative literature on the genus.

A revision of this genus occurring in Ghana is warranted because of the monotonous concentration of research on *B. sapida* (Omobuwajo et al., 2000; Abolaji et al., 2007; Sinmisola et al., 2019; Akintola et al., 2020) and the knowledge gap from Hutchinson & Dalziel (1963). In addition, POWO (2024), recognises 7 and 10 synonyms for *B. sapida* and *B. welwitschii* respectively.

The objective of the study is to update the taxonomy of members in *Blighia* occurring in Ghana. Specifically, construct an identification key for the members of the genus; describe the members of the genus occurring in Ghana, and generate distribution maps of the members of the genus occurring in Ghana.

## METHODOLOGY

Both fieldwork and herbarium studies were conducted for this study. The 2 available fossil records were examined together with 40 herbarium specimens, 20 digital collections from GBIF, and 20 live samples of *Blighia*. 5 morphological characters (leaf length, leaf width, number of secondary veins, length of secondary veins, and length of intersection between the secondary veins) were measured for the fossils. 20 morphological characters (leaf length, leaf width, primary midrib, number of secondary veins, length of each secondary veins, intersection between secondary veins, fruit length, fruit width, leaf arrangement, leaf shape, leaf apex, leaf base, leaf margin, venation type, leaf branching type, leaf surface, inflorescence type, fruit type, seed colour and shape) were measured using herbarium voucher specimens, digital collections and live specimens. Leaves with some parts covered, bent, or broken off were exempted from the measurements. The live specimens were collected from 8 different localities across the country. Collecting *Blighia sapida* from Aburi, Kwahu, Kyebi, Legon, Achimota, Nsawam, and Konongo and *B. welwitschii* from Legon, Achimota, Atewa forest reserve, and Aburi botanical gardens. The period for collection was from January to April. Georeferencing software: GeoPick v.2.1.0, Google Earth Pro v.7.3.6, Google Maps v.6.112.2, and GeoNames v.2.0 were used to validate the accuracy of coordinates provided on the labels of the voucher specimens examined. The validated coordinates were then used to draw distribution maps using SimpleMappr (http://www.simplemappr.net).

## RESULTS AND DISCUSSION

### Identification key

The genus consists of trees; leaves large, even-pinnate; producing fruits with fleshy 3-valved capsules encapsulating seeds large, black to brown and whitish to yellowish fleshy aril.

1a. Leaf oblanceolate sometimes obovate, apex rounded sometimes emarginate, base acute, midrib 5-12 cm long, venation compound rectipinnate… ***B. sapida***

1b. Leaf lanceolate, attenuate sometimes acute, base cuneate, midrib 6-20 long, venation pinnate… ***B. welwitschii***

### Taxonomic treatment

1. *Blighia sapida* K.D.Koenig in Ann. Bot. (König & Sims) 2: 571 (1806)

#### Type

India, Wight, R. 386 (Typus K! K000701709).

#### Synonyms

*Cupania sapida* (K.D.Koenig) Oken in Allg. Naturgesch. 3(2): 1337 (1841); *Akea solitaria* Stokes in Bot. Mat. Med. 2: 354 (1812); *Akeesia africana* Tussac in Fl. Antill. 1: 66 (1808); *Bonannia nitida* Raf. in Specchio Sci. 15: 116 (1814); *Cupania akeesia* Cambess. ex Spach in Hist. Nat. Vég. 3: 60 (1834); *Cupania edulis* Schumach. & Thonn. in C.F.Schumacher, Beskr. Guin. Pl.: 190 (1827); *Sapindus obovatus* Wight & Arn. in Prodr. Fl. Ind. Orient. 1: 111 (1834) (POWO, 2024).

#### Morphological description

***Tree*** height 6-32m high with young branchlets and spreading crown; leaves, 5-15 cm long, 3-8 cm wide. ***Leaf*** opposite and distichous, oblanceolate or obovate, rounded or emarginate at the apex, acute at the base, entire, venation compound rectipinnate, petiole porrect. Primary vein 5–12 cm long. Secondary veins 13–20 measuring up to 6 cm long, intersection between secondary veins up to 4 cm long. Leaflets 3–5 pairs, opposite or subopposite, glabrous above and beneath. ***Fruit*** scarlet, glabrate, 1-12 cm long, aril edible when ripe. ***Seed*** black, oblong 2-5 cm long, 1-3 cm wide, each with a large mustard aril; aril 1-3 cm long, 1.5-3cm wide.

#### Distribution

Kpedze (Volta region), Achimota (Greater Accra region), Kumasi (Ashanti region), Begoro, (Eastern region), Abor-Anyako (Eastern region), Aiyaola Forest Reserve, (Eastern region), Legon Botanical Gardens (Greater Accra region), Atebubu-Amanfrom (Brong-Ahafo region).

#### Habitat and Ecology

The species is distributed across semi-deciduous and evergreen forests, as well as forest outliers in savanna areas. The widespread planting of surrounding communities and extending into the forest further obscures the species’ native habitat. It grows best in deep, fertile soils with good drainage, but it can also be found on limestone and sandy, infertile soil. (PROTA, 2024)

#### Conservation status

Least Concern (IUCN, 2019).

#### Phenology

Flowers throughout the year, mostly in March and April (GBIF, 2023).

#### Vernacular names

Ankye (Twi), Anke (Twi) (Osei et al., 2014).

#### Ethnobotanical uses

The wood of *Blighia sapida* serves as a ‘niangon’ alternative in Ghana. In addition, the wood is utilized to make charcoal and firewood. It is customary to plant *Blighia sapida* as a decorative shade tree. It is thought to be beneficial for controlling erosion and improving soil. In traditional medicine, conjunctivitis and ophthalmia are treated by injecting sap from terminal buds into the eyes (Burkill, 2000). To cure oedema, intercostal pain, dysentery, and diarrhea, decoctions of bark and leaves are used. In Ghana, the body is stimulated by bark mixed up with capsicum pepper (*Capsicum annuum L*.), and migraines are treated by rubbing the pulp of ground leafy twigs on the forehead. The Krobo people of Ghana use green fruits to lather in water and as a dye-making mordant and soap. Potash-rich dried fruit husks are used to make soap from their ashes (PROTA, 2024).

#### Specimens examined

Volta region: Kpedze, 1 January 1925, *F. N. Howes* 1116 (GC); Kpedze, 25 January 1926, *F. N. Howes* 1116 (GC); Greater Accra region: Achimota, 1 May 1931, *F. R. Irvine* 1637; Achimota, 24 June 1935 G. K. Akpabla 297 (GC); University of Ghana Botanical Gardens [05°39’ 36’’, 00°11’11’’W], 20 March 1994, *A. Welsing, N. Merello & H. Schmidt* 13 (GC); Ashanti region: Kumasi, 1 July 1953, Addo-Ashong; Kumasi, 1 July 1954, *Addo-Ashong F. W* (GC); Eastern region: Arboretum, 4 April 1954, *N. F. Madelin; Begoro*, 24 February 1956, *J. K. Morton* A1838; Abor-Anyako Rd, 14 November 1957, *A. A. Enti* 598 (GC); Aiyaola Forest Reserve, Near Kade A. R. S. [6°47’N, 0°54’W], 25 March 1994, *C. C*.*H. Jongkind, D. K. Abbiw & C. M. J. Nieuwenhuis* 1377 (GC); Brong-Ahafo Region: Ca. 7.0 Km South of Atebubu, Along The Atebubu-Amanfrom Road [07°43’ 08’’N, 00°59’ 10’’W], 30 March 1996, *H. H. Schmidt, M. Merello & A. Welsing* 2123 (GC).

2. *Blighia welwitschii* (Hiern) Radlk. in H.G.A.Engler (ed.), Pflanzenr., IV, 165: 1146 (1933)

#### Type

Congo, the Democratic Republic, M. Laurent 940 (Typus BR! BR0000008977762)

#### Synonyms

*Phialodiscus welwitschii* Hiern in Cat. Afr. Pl. 1: 171 (1896); *Blighia kamerunensis* Radlk. in H.G.A.Engler (ed.), Pflanzenr., IV, 165: 1145 (1933); *Blighia laurentii* De Wild. in Ann. Mus. Congo Belge, Bot., sér. 5, 3: 113 (1909); Blighia mildbraedii Radlk. in G.W.J.Mildbraed (ed.), Wiss. Erg. Deut. Zentr.-Afr. Exped., Bot. 2: 480 (1912); *Blighia welwitschii* var. *bancoensis* (Aubrév. & Pellegr.) Pellegr. in Mém. Publiés Soc. Bot. France 1955: 62 (1956); *Blighia wildemaniana* Gilg ex Radlk. in H.G.A.Engler (ed.), Pflanzenr., IV, 165: 1145 (1933); *Phialodiscus bancoensis* Aubrév. & Pellegr. in Bull. Soc. Bot. France 85: 291 (1938); *Phialodiscus laurentii* (De Wild.) Radlk. in H.G.A.Engler (ed.), Pflanzenr., IV, 165: 1149 (1933), nom. illeg.; *Phialodiscus mortehanii* De Wild. in Bull. Jard. Bot. État Bruxelles 4: 361 (1914) (POWO 2024).

#### Morphological description

***Tree***, 7-36m high, with cylindrical trunk; with grey bark showing small reticulate fissures; pistachio mottled. ***Leaf*** opposite and distichous, lanceolate, bluntly attenuate or acute at the apex, sometimes cuneate at the base, entire, venation pinnate, glabrous above and beneath. Petiole porrect. Midrib impressed above, prominent beneath. Leaflets in 2–4 pairs, 5-25 cm long, 3-10 cm. wide, primary veinlet 6-20 cm long. Secondary veinlets measuring up to 6 cm long. ***Fruit*** scarlet, 3-7 cm long, 5-10 cm wide. ***Seeds*** black or bistre, oblong, 2.4-3 cm long, 1.5-6 cm wide; mustard aril 1-3 cm long, 1.5-3cm wide.

#### Distribution

Akim-Manso (Eastern Region), Ateiku (Western Region), Atewa forest reserve (Eastern Region), Aburi botanical garden (Eastern Region).

#### Habitat and Ecology

Reported in closed forests, primary and secondary formations, and on plateaus at 450 to 600 m of altitude (Tropical timbers, 2024)

#### Conservation status

*Blighia welwitschii* is listed as Least Concern. (IUCN Red List, 2018).

#### Phenology

Occurs throughout the year.

#### Vernacular names

Akee (English) (PROTA, 2024).

#### Ethnobotanical uses

Enema with bark mashed and diluted in water (PROTA, 2024).

#### Specimens examined

Eastern region: Manso, Akim, Dec. 1933, *F. R. Irvine*, 2094; Eastern region: Atewa range F. R., 4 Feb. 1971, *J. B. Hall*, 42502 (GC); Western region: Ateiku, 1 Nov. 1973, *Hall and Swaine*, 44593 (GC); Eastern region: Atewa range F. R., 14 Feb 1974, *Hall and Swaine*, 44781 (GC); Eastern region: Aburi botanical garden, 6 Mar. 1976, *Hall, Lock and Abbiw*, 45863 (GC).

*Distribution maps*

**Figure 1.**
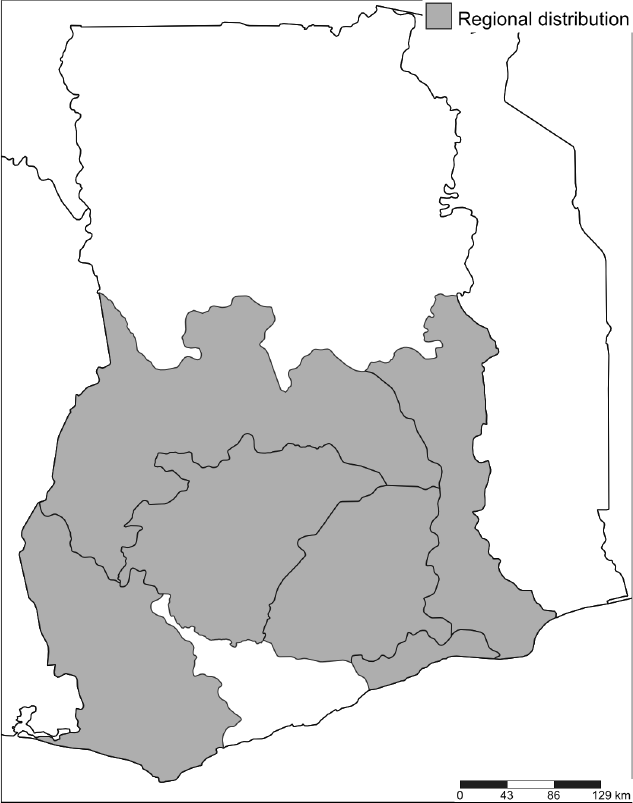
Regional distribution of *Blighia* in Ghana

**Figure 2.**
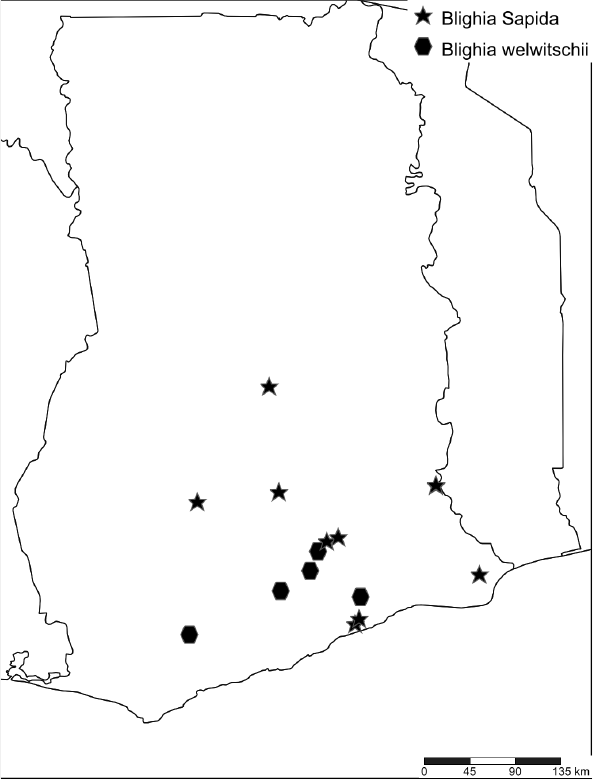
: Species distribution of *Blighia* in Ghana

## ACKNOWLEDGEMENTS

Authors are grateful to the curators in the Ghana Herbarium (GC) and Professor Gabriel K. Ameka for logistics support. Also, Christiana Peprah Oppong for useful comments on the manuscript.

